# The in vivo inhibitory function of the MHC-I α3 domain–CD8α interaction

**DOI:** 10.64898/2026.05.21.726907

**Authors:** Juanjuan Zhao, Lawrence Feng, Yang Xu, Kareem Alsbei, Jun Wang, Tiffany Zhou, Qipeng Zhan, Shiya Sun, Eric Hong, Lingbin Meng, Ning Jin, Xiaolin Cheng, Haitao Wen, Gang Xin, Mark Rubinstein, Stanley Huang, Zihai Li, Xue Han, Linghua Zheng

## Abstract

The interaction between the Major Histocompatibility Complex Class I (MHC-I) α3 domain and CD8α has classically been viewed as a positive coreceptor interaction that stabilizes TCR signaling during antigen recognition. However, its physiological function in mature peripheral CD8+ T cells in vivo remains incompletely understood. Here, we identify the MHC-I α3 domain–CD8α interaction as a previously unrecognized inhibitory pathway that tonically restrains peripheral CD8+ T-cell activation and maintains T-cell tolerance in vivo. Antibody-mediated disruption of the MHC-I α3 domain–CD8α interaction induced spontaneous activation of peripheral CD8+ T cells without impairing their survival, lowered the threshold for antigen-induced activation, and enhanced responsiveness to cognate peptide stimulation. In peptide-induced OT-I T-cell anergy models, blockade of either H-2D^b^ or H-2K^b^ α3 domain interactions with CD8α prevented the induction of anergy and restored responsiveness of previously anergic T cells. Notably, blockade of the H-2K^b^ α3 domain enhanced OT-I responses despite simultaneously disrupting the classical positive coreceptor interaction within the TCR–peptide–MHC complex, indicating that tonic inhibitory signaling mediated by the MHC-I α3 domain predominates under these conditions. Together, these findings redefine the classical MHC-I–CD8α interaction as a bidirectional pathway that not only supports antigen recognition but also imposes tonic inhibitory control over peripheral CD8+ T cells. These results identify the MHC-I α3 domain–CD8α axis as a potential target for reversing T-cell tolerance and enhancing antitumor or antiviral immunity.

## Introduction

The interaction between the Major Histocompatibility Complex Class I (MHC-I) α3 domain and CD8α was identified nearly 40 years ago (*1, 2*), and is classically viewed as a positive coreceptor interaction that stabilizes the TCR–MHC-I complex and augments signaling by low-affinity or CD8-dependent TCRs in vitro (*3-5*). However, these in vitro assays do not fully capture the physiological in vivo functions of the MHC-I α3 domain–CD8α interaction. Although it has been shown that disruption of this interaction profoundly alters positive and negative selection in the thymus (*6, 7*), its physiological function in mature peripheral CD8+ T cells in vivo remains poorly defined.

We recently identified CD8α as an inhibitory checkpoint that maintains peripheral CD8+ T-cell quiescence through interaction with the myeloid-restricted ligand PILRα (*8*). In contrast to PILRα, MHC-I is constitutively expressed by nearly all nucleated cells (*9*), raising the possibility that tonic engagement of CD8α by the MHC-I α3 domain provides a more pervasive inhibitory signal under homeostatic conditions. We therefore hypothesized that continuous interaction between the MHC-I α3 domain and CD8α restrains spontaneous CD8+ T-cell activation in the periphery and establishes the threshold for antigen responsiveness.

Here, we show that in vivo antibody-mediated disruption of the MHC-I α3 domain–CD8α interaction induces spontaneous activation of peripheral CD8+ T cells without impairing their survival, lowers the threshold for antigen-induced activation, prevents the induction of T-cell anergy, and reverses established anergy in vivo. These findings identify the MHC-I α3 domain–CD8α axis as a previously unrecognized inhibitory pathway that tonically restrains CD8+ T-cell activity and maintains peripheral tolerance.

## Results

### Tonic MHC-I α3 domain–CD8α interaction restrains spontaneous CD8^+^ T-cell activation in vivo

To directly examine the in vivo function of the MHC-I α3 domain–CD8α interaction, we used mAb clone 28-14-8S, which recognizes the α3 domain of both H-2D^b^ (*10*) and H-2L^d^ (*11*). Flow cytometric analysis confirmed that 28-14-8S effectively disrupted binding between H-2L^d^ and CD8α (**Fig. 1A**).

**Fig. 1.**
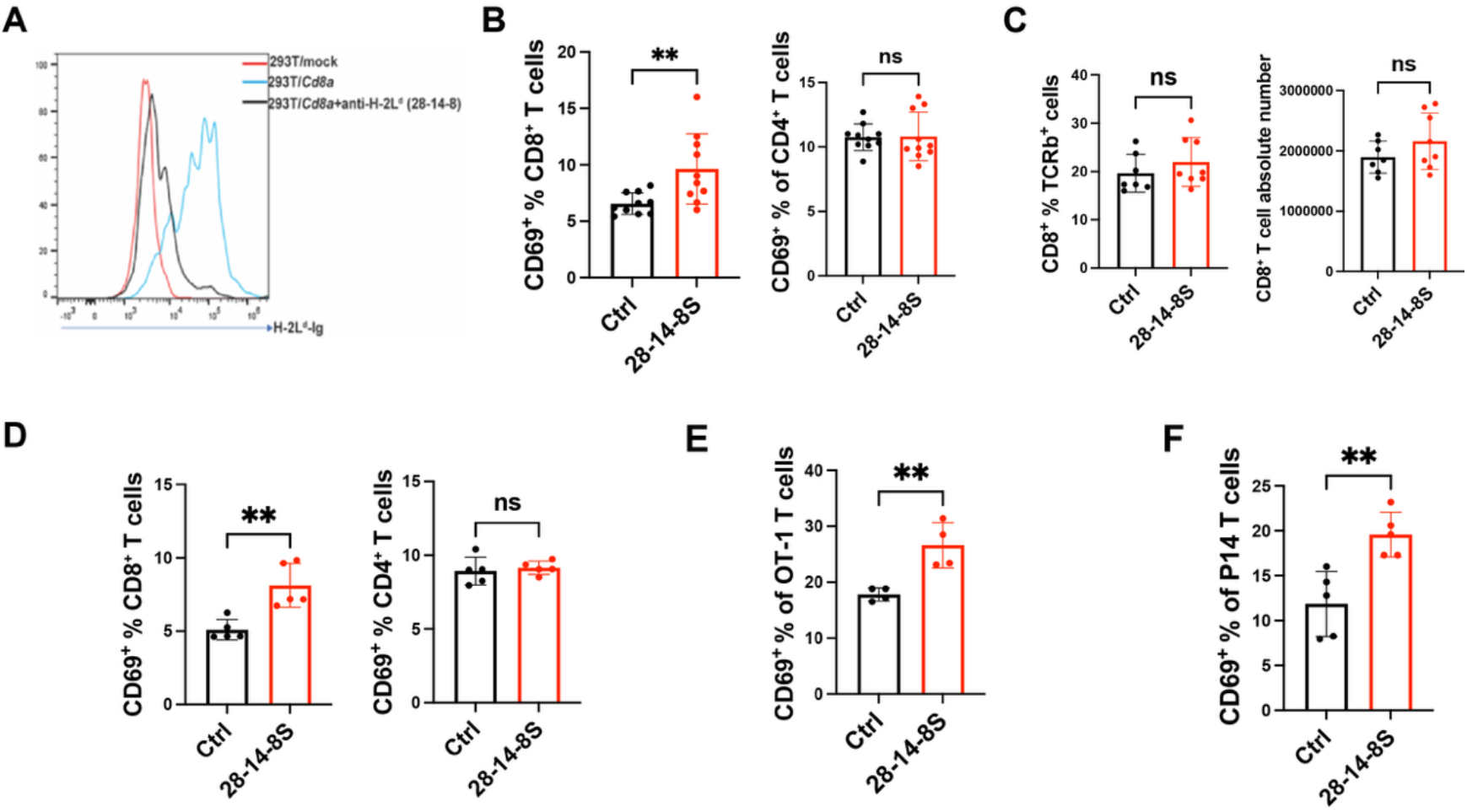
Blockade of the MHC-I α3 domain–CD8α interaction disrupts CD8^+^ T cell quiescence. **(A)**Flow cytometric analysis of H-2L^d^-Ig binding to mock- or CD8α-transfected 293T cells in the presence or absence of anti-H-2L^d^ monoclonal antibody (mAb) 28-14-8S. **(B)**CD69 expression on lymph node CD8^+^ and CD4^+^ T cells 24 h after in vivo treatment of BALB/c mice with 28-14-8S, which blocks the interaction between the H-2L^d^ α3 domain and CD8α(n=10 mice per group). **(C)**In BALB/c mice, lymph nodes were harvested 12 days after repeated in vivo administration of 28-14-8S (twice weekly). The proportion of CD8^+^ T cells within the TCRβ^+^ population and the absolute number of CD8^+^ T cells in lymph nodes were determined by flow cytometry. **(D)**CD69 expression on lymph node CD8^+^ and CD4^+^ T cells 24 h after in vivo treatment of C57BL/6 mice with 28-14-8S, which blocks the interaction between the H-2D^b^ α3 domain and CD8α. (**E, F**) CD69 expression on OT-I (E) or P14 (F) T cells in the lymph nodes of C57BL/6 mice 24 h after adoptive transfer of OT-I or P14 T cells followed by in vivo treatment with control antibody or 28-14-8S. Data shown in (B to F) are representative of at least two independent experiments. n=10 mice per group in (B); n=7-8 mice per group in (C). n=4-5 mice per group in (D to F). Values represent mean ± SD. P-value was determined by unpaired two-tailed Student’s t-test. ****P < 0.0001, ***P < 0.001, **P < 0.01, *P < 0.05, ns: not significant.

Systemic administration of 28-14-8S in BALB/c mice induced spontaneous activation of peripheral CD8^+^ T cells but not CD4^+^ T cells, as indicated by increased CD69 expression in the absence of exogenous antigen (**Fig.1B**). Unlike blockade of the PILRα–CD8α pathway, which can activate CD8+ T cells and reduce their survival (*8*), disruption of the MHC-I α3 domain–CD8α interaction did not reduce CD8+ T-cell survival 12 days after antibody treatment (**Fig. 1C**).

A similar increase in spontaneous CD8^+^ T-cell activation but not CD4 T-cell activation was observed in vivo in C57BL/6 mice following treatment with 28-14-8S (**Fig. 1, D**). Thus, blockade of either H-2L^d^ in BALB/c mice or H-2D^b^ in C57BL/6 mice was sufficient to induce spontaneous CD8^+^ T-cell activation, suggesting that tonic inhibition is not unique to a single MHC-I molecule. Because CD8^+^ T cells in C57BL/6 mice are predominantly restricted by either H-2K^b^ or H-2D^b^, we next asked whether blockade of the H-2D^b^ α3 domain selectively affected only the H-2K^b^ or H-2D^b^-restricted subset. OT-I (H-2K^b^-restricted) or P14 (H-2D^b^-restricted) transgenic CD8^+^ T cells were therefore transferred into wild-type C57BL/6 recipients, followed by antibody treatment. In vivo blockade of the H-2D^b^ α3 domain activated both OT-I and P14 cells (**Fig. 1, E** and **F**), indicating that tonic MHC-I α3 domain–CD8α-mediated inhibition is not confined to a single MHC-I-restricted CD8^+^ T-cell population.

To exclude clone-specific effects, we examined two additional mAbs, clones 34-2-12 and SF1-1.1.10, which have been reported to specifically recognize the α3 domain of H-2D^d^ and H-2K^d^, respectively (*12-14*). Both antibodies disrupted the binding of the MHC-I α3 domain to CD8α (**fig. S1A and B**) and similarly induced antigen-independent activation of CD8^+^ T cells in BALB/c mice (**fig. S1C and D**). Together, these findings indicate that the MHC-I α3 domain–CD8α interaction functions as a tonic inhibitory rheostat that restrains spontaneous CD8^+^ T-cell activation under homeostatic conditions.

### In vivo disruption of the MHC-I α3 domain–CD8α interaction lowers the threshold for CD8^+^ T cell activation and enhances antigen responsiveness

We next investigated whether tonic inhibition mediated by the MHC-I α3 domain–CD8α axis regulates the activation threshold and antigen responsiveness of CD8^+^ T cells. C57BL/6 mice express two classical MHC-I molecules, H-2D^b^ and H-2K^b^. To interrogate the MHC-I α3 domain–CD8α axis comprehensively, we generated a novel anti–H-2K^b^ α3 domain monoclonal antibody, clone 3C11, which disrupts the H-2K^b^–CD8α interaction. C57BL/6 mice were treated in vivo with anti–H-2D^b^ α3 domain mAb 28-14-8S or anti–H-2K^b^ α3 domain mAb 3C11. Splenocytes were subsequently harvested and stimulated ex vivo with PMA plus ionomycin to assess global activation responsiveness independent of TCR specificity. H-2D^b^ or H-2K^b^ α3 domain blockade significantly increased CD25 expression in CD8^+^ T cells (**Fig. 2A**), suggesting that disruption of the MHC-I α3 domain–CD8α interaction lowers the activation threshold of peripheral CD8^+^ T cells.

**Fig. 2.**
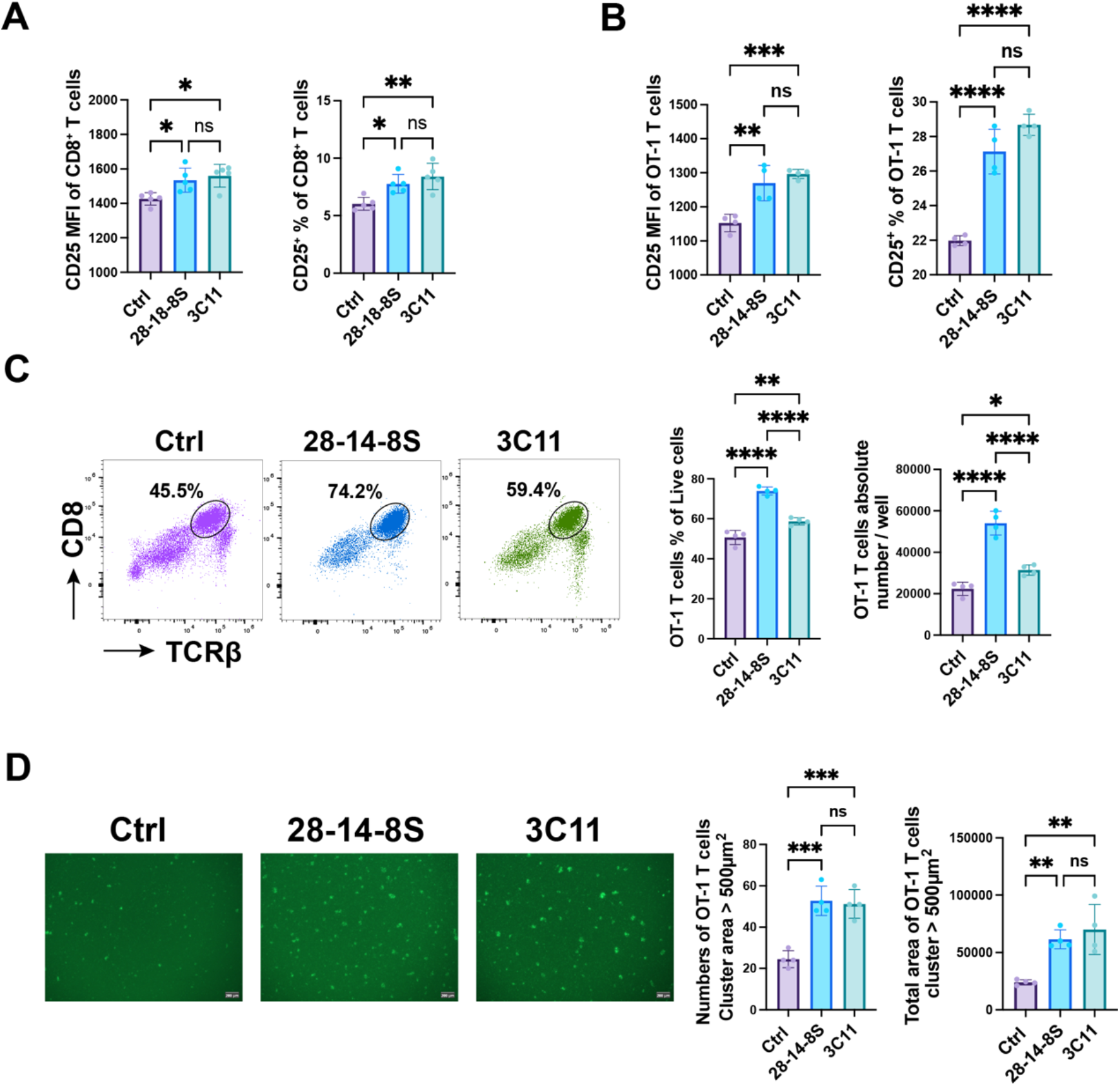
MHC-I α3 domain–blocking antibodies lower the activation threshold of CD8^+^ T cells. (**A**) C57BL/6 mice were euthanized 24 h after in vivo administration of anti-H-2Db mAb 28-14-8S or anti-H-2Kb mAb 3C11. Splenocytes were subsequently stimulated in vitro with PMA and ionomycin for 6 h, followed by analysis of CD25 expression on CD8^+^ T cells. (**B, C**) OT-I mice were treated in vivo with MHC-I α3 domain–blocking antibody (28-14-8S or 3C11). Twenty-four hours later, splenocytes were stimulated in vitro with OVA peptide (10 ng/mL). CD25 expression on OT-I T cells was analyzed 6 h after stimulation (B). After 48 h, both the 28-14-8S and 3C11 treatment groups showed significantly increased frequencies and absolute numbers of CD8^+^ T cells compared with the control group (Ctrl) (C). (**D**) OT-I mice were treated in vivo with 28-14-8S, 3C11, or Ctrl and euthanized 24 h later. OT-I T cells were isolated, labeled with CFSE, and co-cultured with splenocytes from wild-type C57BL/6 mice at a 1:3 ratio in the presence of OVA peptide (10 ng/mL). After 24 h of co-culture, fluorescence microscopy revealed significantly increased numbers of OT-I T cell clusters with areas >500 μm^2^, as well as greater total cluster area, in the 28-14-8S and 3C11 treatment groups compared with Ctrl. n=5 mice per group in (A); n=4 mice per group in (B). Values represent mean ± SD. P value was determined by One-Way ANOVA (Analysis of Variance). ****P < 0.0001, ***P < 0.001, **P < 0.01, *P < 0.05, ns: not significant.

To determine whether this effect also enhances antigen-specific responses, OT-I transgenic mice were treated in vivo with 28-14-8S or 3C11 and splenocytes were stimulated ex vivo with SIINFEKL peptide. Blockade of the MHC-I α3 domain significantly enhanced OT-I sensitivity to peptide stimulation, as evidenced by increased CD25 expression (**Fig. 2B**). Notably, blockade of the H-2K^b^ α3 domain with 3C11 antibody also resulted in elevated CD25 expression on OT-1 T cells, even though the OT-I TCR is H-2K^b^-restricted(*15*). In addition, treatment with either 28-14-8S or 3C11 increased the number of OT-I cells recovered after peptide stimulation (**Fig. 2, C and D**). Thus, in vivo disruption of either the H-2D^b^ or H-2K^b^ α3 domain–CD8α interaction enhances subsequent OT-I activation in response to cognate peptide stimulation in vitro.

### In vivo disruption of the MHC-I α3 domain–CD8α interaction prevents the induction of CD8^+^ T cell anergy

Because this pathway regulates activation thresholds, we next tested whether it also contributes to peripheral tolerance induction. In a peptide-induced CD8^+^ T cell anergy model (*16, 17*), in vivo administration of either 28-14-8S or 3C11 during priming enhanced OT-I CD8^+^ T cell responses and prevented the induction of anergy (**Fig. 3, A-C**). In control-treated mice, peptide rechallenge on day 31 failed to induce expansion of OT-I cells in the blood, consistent with successful anergy induction. In contrast, in mice treated with 28-14-8S or 3C11, peptide rechallenge induced robust OT-I proliferation, with significantly increased frequencies of OT-I CD8+ T cells detected in the blood on days 35, 39, and 42 (**Fig. 3B, C**). Thus, blockade of either the H-2D^b^ α3 domain–CD8α interaction or the H-2K^b^ α3 domain–CD8α interaction during priming enhances OT-I expansion and prevents the subsequent development of anergy in this model.

**Fig. 3.**
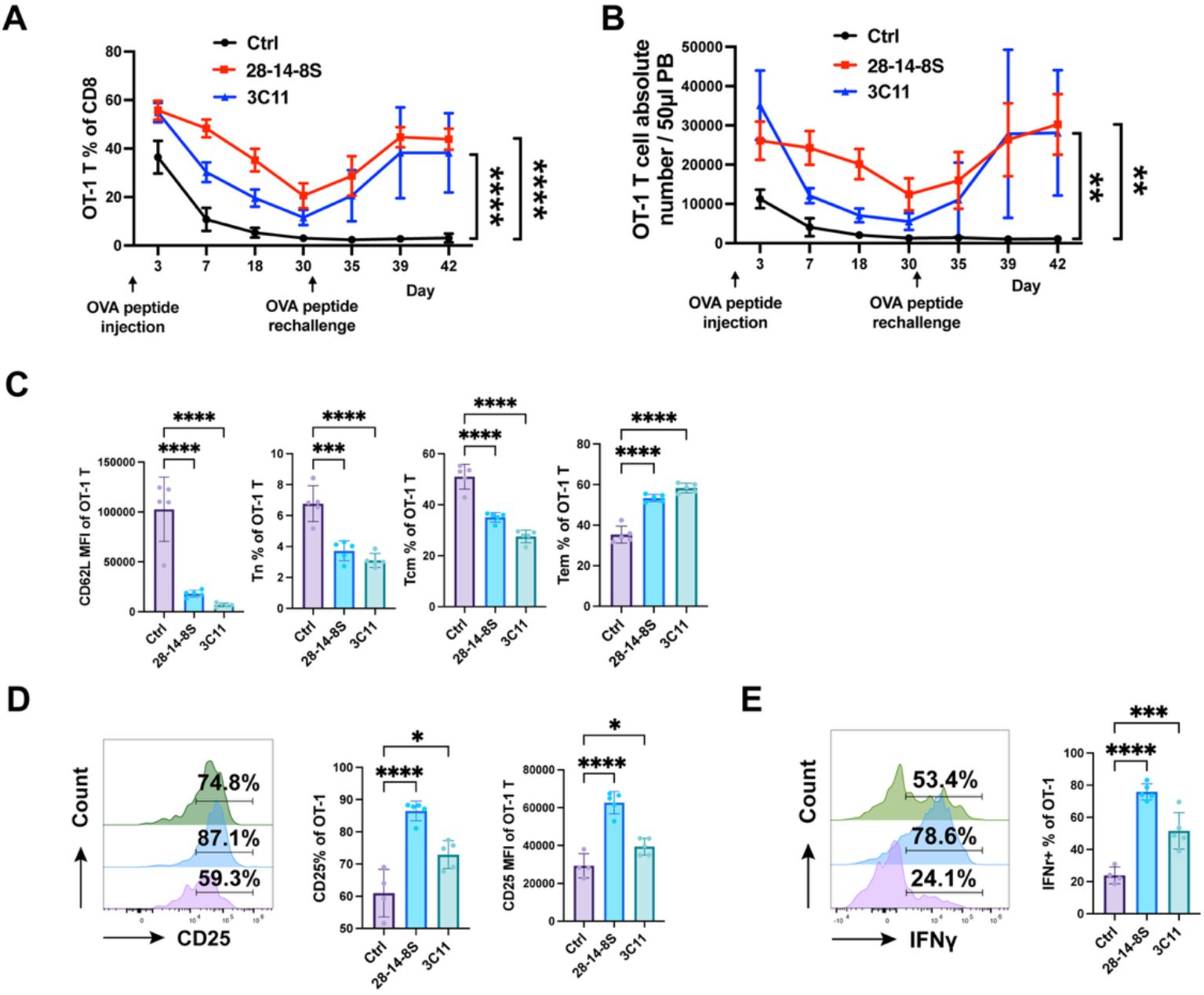
MHC-I α3 domain–blocking antibodies prevent OT-I T cell anergy. (**A** to **C**) Ova peptide was given (i.v) on days 0 and 31. Antibodies were given (i.p.) on day 0. Kinetic changes in the proportion of OT-I T cells among peripheral blood CD8^+^ T cells were detected by flow cytometry (A). Kinetic changes in the absolute number of OT-I T cells per 50 μL of peripheral blood (PB) was detected by flow cytometry (B). On day 3, expression of the differentiation-associated marker CD62L on blood OT-I T cells, as well as the proportions of naïve T cells (T_n; CD62L^+^CD44^−^), central memory T cells (T_cm; CD62L^+^CD44^+^), and effector memory T cells (T_em; CD62L^−^CD44^+^), were analyzed by flow cytometry (C). (**D** and **E**) Ova peptide was given (i.v) on day 0, and antibodies were given (i.p.) on day 0. On day 7, mice were euthanized, and splenocytes were subsequently stimulated in vitro with OVA peptide (10 ng/mL). Expression of CD25 (D) and IFN-γ (E) in splenic OT-I T cells was detected by flow cytometry. Data shown in (A to E) are representative of two independent experiments. Each symbol represents one mouse. n=5 mice per group in (A to C); n=4-5 mice per group in (D and E). Values represent mean ± SD. P value was determined by One-Way ANOVA (Analysis of Variance). ****P < 0.0001, ***P < 0.001, **P < 0.01, *P < 0.05, ns: not significant.

Although **Fig. 3B and C** show the increase in circulating OT-I cells, similar increases were also observed in the lymph nodes and spleen, indicating that the effect was systemic rather than restricted to the blood (**fig. S2A–D**).

Before peptide rechallenge on day 31, during the anergy-induction phase, in addition to OT-I cell number, the activation status was also altered by antibody treatment. OT-I cells treated with 28-14-8S or 3C11 showed reduced CD62L expression and an increased proportion of CD44^+^CD62L^-^ OT-I cells (**Fig. 3D**). In addition, CD25 expression (**Fig. 3E**) and IFN-γ production (**Fig. 3F**) in OT-I cells were increased in both treatment groups. These findings indicate that disruption of the MHC-I α3 domain–CD8α interaction promotes early OT-I activation and opposes the establishment of the anergic state.

Unlike 28-14-8S, which targets H-2D^b^ and therefore cannot directly interfere with recognition of the OT-I-restricting H-2K^b^–peptide complex, 3C11 disrupts the interaction between CD8α and the restricting H-2K^b^ molecule itself. Thus, in this model, 3C11 would be expected to exert two opposing effects simultaneously: inhibition of the classical positive coreceptor function of H-2K^b^–CD8α during antigen recognition, and relief of tonic inhibitory signaling mediated by the H-2K^b^ α3 domain–CD8α interaction. Despite this, 3C11 still enhanced OT-I responses and prevented anergy, indicating that under these conditions the inhibitory function of the H-2K^b^ α3 domain–CD8α interaction predominates over its classical costimulatory role. Together, these results indicate that tonic signaling through the MHC-I α3 domain–CD8α axis contributes to the establishment of peripheral CD8+ T-cell tolerance.

### In vivo blockade of the MHC-I α3 domain–CD8α interaction reverses established T-cell anergy

We next asked whether the MHC-I α3 domain–CD8α interaction is required only for the induction of CD8+ T-cell tolerance or also for its maintenance. Using the same peptide-induced OT-I anergy model, previously anergized OT-I cells were rechallenged in vivo with SIINFEKL peptide in the presence of either 28-14-8S or 3C11. In control-treated mice, OT-I cells remained hyporesponsive to peptide rechallenge in the blood, indicating an anergic status of OT-I cells. By contrast, blockade of the H-2D^b^ α3 domain–CD8α interaction with 28-14-8S or the H-2K^b^ α3 domain–CD8α interaction with 3C11 induced robust proliferation of OT-I cells in the blood at multiple time points after peptide rechallenge (**Fig. 4, A–C**), indicating reversal of established anergy.

**Fig. 4.**
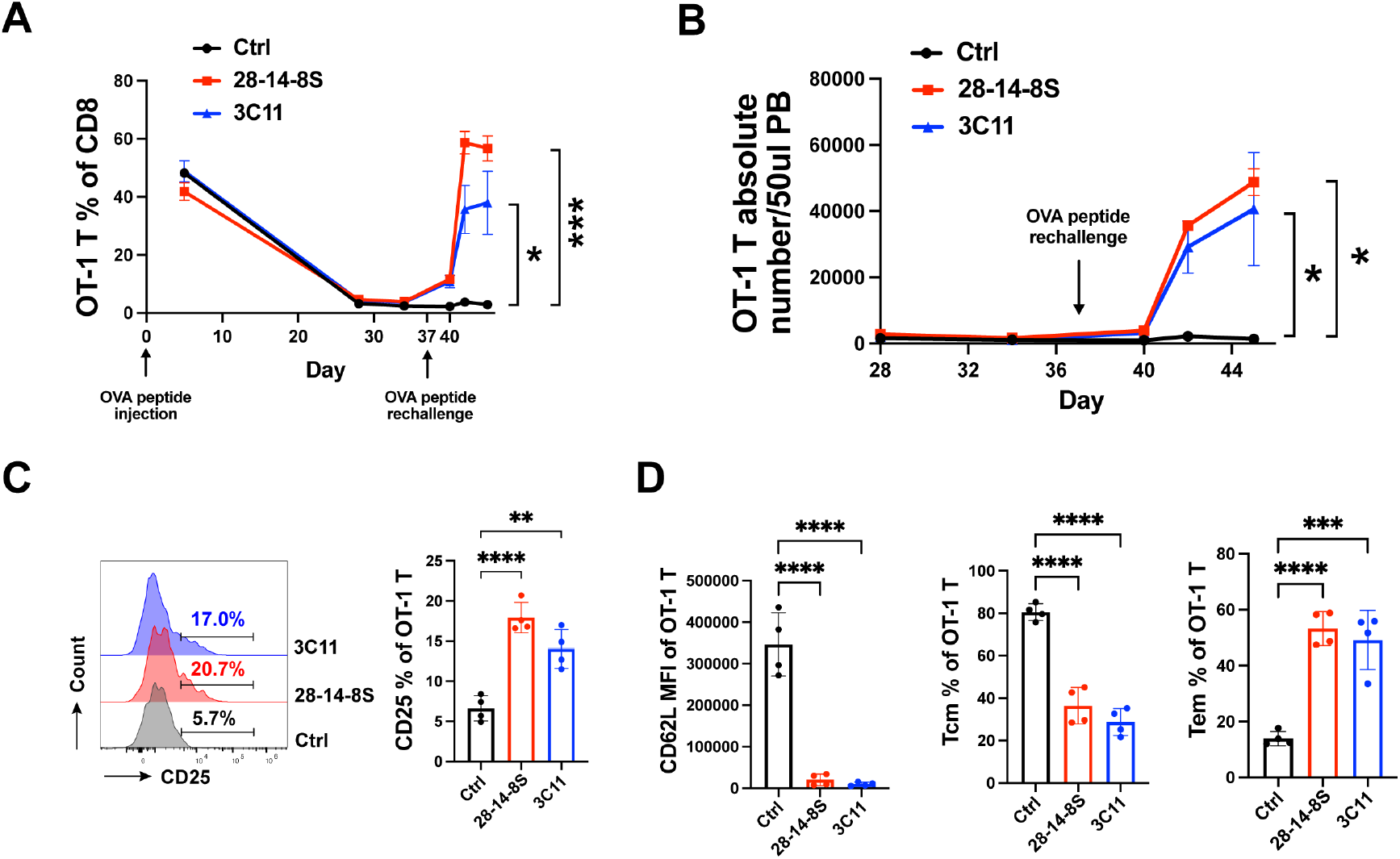
MHC-I α3 domain–blocking antibodies reverse OT-I T cell anergy. OT-I T cells were intravenously (i.v.) transferred on Day −1, and OVA peptide was administered intravenously on Days 0 and 37. Control antibody, 28-14-8S, or 3C11 was administered intraperitoneally (i.p.) on Days 37, 40, and 41. Peripheral blood mononuclear cells (PBMCs) were analyzed on at the indicated time points. **(A)**Kinetic changes in the proportion of OT-I T cells among peripheral blood CD8^+^ T cells. **(B)**Kinetic changes in the absolute number of OT-I T cells per 50 μL of peripheral blood. **(C)**Expression of the activation marker CD25 on OT-I T cells in the blood on day 40 (3 days after peptide rechallenge). **(D)**On day 40 (3 days after peptide rechallenge), expression of the differentiation-associated marker CD62L on OT-I T cells, as well as the proportions of central memory T cells (T_cm_; CD62L^+^CD44^+^) and effector memory T cells (T_em_; CD62L^−^CD44^+^) in the blood, were analyzed by flow cytometry. Data shown are representative of two independent experiments. Each symbol represents one mouse. n=4 mice per group. Values represent mean ± SD. P value was determined by One-Way ANOVA (Analysis of Variance). ****P < 0.0001, ***P < 0.001, **P < 0.01, *P < 0.05, ns: not significant.

In addition to increased OT-I cell numbers, antibody treatment restored an activated phenotype. Three days after peptide rechallenge, OT-I cells from mice treated with 28-14-8SS or 3C11 expressed higher levels of CD25 and lower levels of CD62L, with a greater proportion of CD44^+^CD62L^lo^ cells in the blood (**Fig. 4, D and E**).

A similar restoration of responsiveness was observed in the spleen. Treatment with either 28-14-8SS or 3C11 enabled previously anergic OT-I cells to respond to SIINFEKL rechallenge (**Fig. 5A**). Moreover, following ex vivo peptide stimulation, OT-I cells from the 28-14-8SS– or 3C11-treated groups exhibited significantly increased expression of CD25 and CD69, together with enhanced production of IFN-γ, granzyme B, and TNF-α (**Fig. 5B**). Similar results were obtained when OT-I cells were restimulated ex vivo with B16-OVA cells: OT-I cells from the 28-14-8SS– and 3C11-treated groups displayed increased CD25 and CD69 expression and produced higher amounts of IFN-γ, granzyme B, and TNF-α (**Fig. 5C**).

**Fig. 5.**
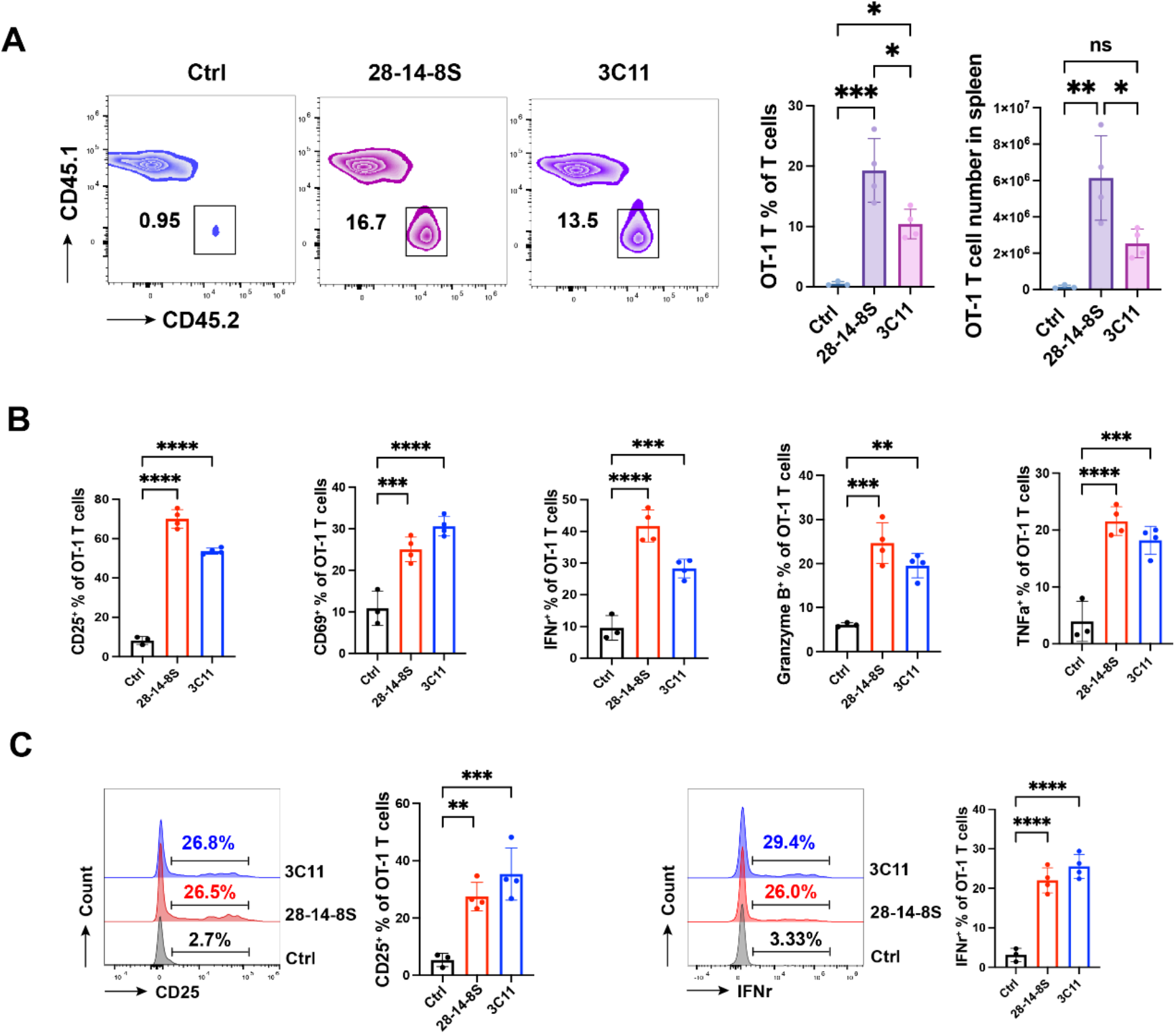
The restoration of responsiveness of OT-1 T cells in the spleen after blockade of MHC-I α3–CD8α interaction. OT-I cells (CD45.2) were transferred into CD45.1 B6 congenic mice, followed by peptide injection (i.v.) to induce anergic OT-I cells. Around 30 days later, the recipient CD45.1 mice were rechallenged with the peptide (i.v.), simultaneously receiving control or 28-14-8S or 3C11 antibody treatment (i.p.). (A)Eight days after OVA peptide rechallenge, the proportions of OT-1 T cells among T cells and the absolute number of OT-1 T cells in the spleen were detected by flow cytometry. (B)Eight days after OVA peptide rechallenge, mice splenocytes were stimulated in vitro with OVA peptide (10 ng/mL). After 24 hours, the expression of the activation markers CD25 and CD69 on OT-1 T cells, as well as the IFN-γ, Granzyme B, and TNF-α in OT-I cells were detected by flow cytometry. (C)Eight days after OVA peptide rechallenge, mice splenocytes were co-cultured in vitro with OVA-overexpressing B16 cells. After 24 hours, the expression of CD25 and IFN-γ in OT-1 T cells was detected by flow cytometry. CD45.2+CD45.1-TCRb+ cells were gated as OT-I cells in flow cytometry analysis. Each symbol represents one mouse. n=3-4 mice per group. Values represent mean ± SD. P value was determined by One-Way ANOVA (Analysis of Variance). ****P < 0.0001, ***P < 0.001, **P < 0.01, *P < 0.05, ns: not significant.

Together, these findings demonstrate that the MHC-I α3 domain–CD8α interaction is required not only for the induction of CD8+ T-cell anergy, but also for the maintenance of the established anergic state.

## Discussion

Our study identifies the MHC-I α3 domain–CD8α interaction as a previously unrecognized inhibitory pathway that tonically restrains peripheral CD8+ T-cell activation in vivo. Although the MHC-I α3 domain has long been viewed as a positive coreceptor-binding site that stabilizes TCR engagement during antigen recognition, our findings reveal an unexpected inhibitory function in vivo under steady-state conditions. Acute in vivo disruption of this interaction induced spontaneous activation of peripheral CD8^+^ T cells, lowered the threshold for antigen-induced activation, and prevented or reversed T-cell anergy. These findings suggest that the MHC-I α3 domain–CD8α interaction exerts distinct functions depending on context: during acute cognate antigen recognition, it stabilizes TCR signaling; whereas under steady-state conditions in the absence of antigen, it provides a tonic inhibitory signal that restrains spontaneous activation. The experiments using the anti-H-2K^b^ α3 domain mAb 3C11 further indicate that these two activities can operate simultaneously. Because 3C11 disrupts the interaction between CD8α and the restricting H-2K^b^ molecule within the OT-I TCR–peptide–MHC complex, it would be predicted to impair the classical stimulatory function of CD8α. However, 3C11 nevertheless enhanced OT-I responses, indicating that relief of tonic MHC-I–CD8α-mediated inhibition outweighs loss of positive coreceptor function under these conditions. Thus, the net effect of MHC-I α3 domain blockade reflects the balance between two opposing functions of the same molecular interaction.

These findings extend our recent work identifying CD8α as an inhibitory checkpoint engaged by PILRα (*8*). However, the MHC-I α3 domain–CD8α axis differs fundamentally from the PILRα–CD8α pathway. PILRα is expressed primarily by myeloid cells, including neutrophils, macrophages, mast cells, and monocytes (*18-20*), whereas MHC-I is constitutively expressed by nearly all nucleated cells. MHC-I may therefore provide a more pervasive inhibitory input that continuously calibrates the activation threshold of CD8^+^ T cells throughout the body. In addition, unlike the disruption of the PILRα–CD8α axis (*8*), blockade of the MHC-I α3 domain–CD8α interaction activated CD8^+^ T cells without impairing their survival. This distinction suggests that different CD8α ligands deliver qualitatively distinct inhibitory signals and supports the concept that CD8α integrates multiple tonic inhibitory inputs with nonredundant functions.

Notably, blockade of the MHC-I α3 domain–CD8α interaction not only induced spontaneous activation of naïve CD8^+^ T cells but also reversed established CD8+ T-cell anergy (**Figs. 4 and 5**). These findings suggest that the MHC-I–CD8α axis provides a continuous tonic inhibitory signal that operates in multiple contexts. In naïve peripheral CD8^+^ T cells, this pathway maintains quiescence under steady-state conditions. In previously antigen-exposed T cells, the same tonic inhibitory signal maintains the hyporesponsive state of anergy. Thus, established anergy may not represent an irreversible fate, but rather a state that requires persistent MHC-I α3 domain–CD8α-mediated restraint.

The ability of MHC-I α3 domain blockade to both prevent and reverse CD8^+^ T-cell anergy suggests that this pathway may represent a therapeutically targetable checkpoint for enhancing antitumor or antiviral immunity. More broadly, our findings redefine the classical MHC-I–CD8α interaction from a purely stimulatory coreceptor mechanism to a bidirectional pathway that also imposes tonic inhibition on CD8^+^ T cells in vivo.

## Materials and Methods

### Mice

C57BL/6, BALB/c, and C57BL/6 CD45.1 congenic mice were purchased from Charles River Laboratories. NZBWF1/J mice and OT-I TCR transgenic mice were obtained from Jackson Laboratory. All animal experiments were approved by the Institutional Animal Care and Use Committee (IACUC) at The Ohio State University.

### Functional Antibody Generation, Production, and Purification

Hybridoma cell lines producing monoclonal antibodies against the MHC-I α3 domain, including 28-14-8S, 34-2-12, and SF1-1.1.10, were purchased from the American Type Culture Collection (ATCC). The hybridoma producing anti-H-2K^b^ α3 domain mAb 3C11 was generated in our laboratory, as described previously (*21*).Briefly, H-2K^b^-Ig (BD Biosciences) and H-2K^b^ tetramers obtained from the National Institutes of Health (NIH) Tetramer Core Facility were used to immunize NZBWF1/J mice, which were subsequently screened for the selected clone. All antibodies (28-14-8S, 34-2-12, SF1-1.1.10, and 3C11) were mouse IgG and were purified using Protein G HP Column (Cytiva HiTrap™). Either mouse IgG purchased from Bio X Cell or an anti-human PD-1H mAb (mIgG) purified with a Protein G HP Column (Cytiva HiTrap™) in our lab was used as the control antibody.

### Flow Cytometry and Cell Counting

Flow cytometry (FCM) was performed on Cytek 3-laser or 5-laser instruments. For the anti-MHC-I α3 domain antibody blockade assay, the H-2L^d^-Ig was purchased from BD Biosciences, and the H-2D^d^ and H-2K^d^ tetramers were obtained from the NIH Tetramer Core Facility.

For the lymph node CD8+ T cell number determination, six lymph nodes (two of inguinal, two of brachial, and two of axillary) were pooled together to isolate total lymph node cells. CD8+ T cell frequency was determined by flow cytometry. CD8+ T cell number = total lymph node cells × CD8+ T cell frequency.

For the OT-I T cell number in the blood determination, the absolute number of OT-1 T cells in peripheral blood (PB) was monitored by FCM using CountBrightTM Absolute Counting Beads (Invitrogen™, C36950).

For the OT-I cluster experiment, OT-I T cell clusters were captured under a fluorescence microscope. The number of cell clusters larger than 500 μm^2^ in a random field was quantified using ImageJ.

### OT-I T Cell Anergy Model

In the prevention of OT-I anergy formation model, C57BL/6 wild-type or C57BL/6 CD45.1 congenic recipient mice were intravenously (i.v) administered 1 million OT-I T cells per mouse (Day −1). OVA peptide (257–264, SIINFEKL, 100 μg per mouse) was administered (i.v) and mice were simultaneously treated (i.p) with either an MHC-I α3 blocking antibody (28-14-8S or 3C11) or the control antibody (Day 0).) Around 30 days later, mice were rechallenged with OVA peptide (100 μg per mouse). And the OT-I cell frequency was detected at the indicated time points.

In the reverse of the established anergic OT-I model, C57BL/6 CD45.1 congenic recipient mice were administered (i.v) 1 million OT-I T cells per mouse (Day −1). OVA peptide (100 μg per mouse) was administered (i.v) (Day 0). The proportion and absolute number of OT-1 T cells in PB were monitored. Around 30 days later, mice were rechallenged with OVA peptide (257–264, SIINFEKL, 100 μg per mouse) and simultaneously treated (i.p) with either an MHC class I α3-blocking antibody (28-14-8S or 3C11) or the control antibody. The frequency and absolute number of OT-1 T cells were detected at the indicated time points.

### T Cell Intracellular Staining

Mice splenocytes were harvested and stimulated in vitro with OVA peptide (10 ng/mL) or OVA-overexpressing B16 cells for 24 hours. Brefeldin A (5 μg/ml, Biolegend) was added in the last 4-6 hours. IFN-γ, Granzyme B, and TNF-α in OT-I cells were detected by flow cytometry.

## Statistical Analysis

Statistical analysis was performed using Graphpad Prism 10.0 (GraphPad Software, Inc., La Jolla, CA, USA). Statistical analyses between the two groups were analyzed by an unpaired two-tailed Student’s t-test. Statistical analyses among the three groups were performed by one-way analysis of variance (ANOVA). Data are presented as mean ± SD, where error bars are shown. P values of less than 0.05 were considered statistically significant. ns, not significant. \**P*<0.05, \*\**P*<0.01, \*\*\**P*<0.001, \*\*\*\**P*<0.0001.

## Author Contributions

J.Z., X.H., and L.Z. designed the study; J.Z., L.F., Y.X., K.A., J.W., T.Z., Q.Z., S.S., and E.H. performed experiments and acquired the data; L.M, N.J, X.C, H.W, G.X, M.R, S.H, and Z.L provided resources and assistance. X.H and L.Z secured funding and supervised the project; X.H and L.Z. conceived and wrote the manuscript.

## References

1. A. M. Norment, R. D. Salter, P. Parham, V. H. Engelhard, D. R. Littman, Cell-cell adhesion mediated by CD8 and MHC class I molecules. Nature 336, 79–81 (1988).

2. J. M. Connolly, T. H. Hansen, A. L. Ingold, T. A. Potter, Recognition by CD8 on cytotoxic T lymphocytes is ablated by several substitutions in the class I alpha 3 domain: CD8 and the T-cell receptor recognize the same class I molecule. Proc Natl Acad Sci U S A 87, 2137–2141 (1990).

3. J. R. Wyer et al., T cell receptor and coreceptor CD8 alphaalpha bind peptide-MHC independently and with distinct kinetics. Immunity 10, 219–225 (1999).

4. K. C. Garcia et al., CD8 enhances formation of stable T-cell receptor/MHC class I molecule complexes. Nature 384, 577–581 (1996).

5. L. Wooldridge et al., Interaction between the CD8 coreceptor and major histocompatibility complex class I stabilizes T cell receptor-antigen complexes at the cell surface. J Biol Chem 280, 27491–27501 (2005).

6. C. J. Aldrich et al., Negative and positive selection of antigen-specific cytotoxic T lymphocytes a_ected by the alpha 3 domain of MHC I molecules. Nature 352, 718–721 (1991).

7. N. Killeen, A. Moriarty, H. S. Teh, D. R. Littman, Requirement for CD8-major histocompatibility complex class I interaction in positive and negative selection of developing T cells. J Exp Med 176, 89–97 (1992).

8. L. Zheng et al., The CD8alpha-PILRalpha interaction maintains CD8(+) T cell quiescence. Science 376, 996–1001 (2022).

9. A. S. Daar, S. V. Fuggle, J. W. Fabre, A. Ting, P. J. Morris, The detailed distribution of MHC Class II antigens in normal human organs. Transplantation 38, 293–298 (1984).

10. H. Allen, D. Wraith, P. Pala, B. Askonas, R. A. Flavell, Domain interactions of H-2 class I antigens alter cytotoxic T-cell recognition sites. Nature 309, 279–281 (1984).

11. G. A. Evans, D. H. Margulies, B. Shykind, J. G. Seidman, K. Ozato, Exon shu_ling: mapping polymorphic determinants on hybrid mouse transplantation antigens. Nature 300, 755–757 (1982).

12. K. Ozato, N. M. Mayer, D. H. Sachs, Monoclonal antibodies to mouse major histocompatibility complex antigens. Transplantation 34, 113–120 (1982).

13. J. P. Abastado, A. Casrouge, P. Kourilsky, Di_erential role of conserved and polymorphic residues of the binding groove of MHC class I molecules in the selection of peptides. J Immunol 151, 3569–3575 (1993).

14. G. Noun et al., Alloreactive monoclonal antibodies select Kd molecules with di_erent peptide profiles. J Immunol 157, 2455–2461 (1996).

15. S. R. Clarke et al., Characterization of the ovalbumin-specific TCR transgenic line OT-I: MHC elements for positive and negative selection. Immunol Cell Biol 78, 110–117 (2000).

16. F. Tsushima et al., Interaction between B7-H1 and PD-1 determines initiation and reversal of T-cell anergy. Blood 110, 180–185 (2007).

17. J. J. Park et al., B7-H1/CD80 interaction is required for the induction and maintenance of peripheral T-cell tolerance. Blood 116, 1291–1298 (2010).

18. Y. Sun et al., PILRalpha negatively regulates mouse inflammatory arthritis. J Immunol 193, 860–870 (2014).

19. M. Kohyama et al., Monocyte infiltration into obese and fibrilized tissues is regulated by PILRalpha.Eur J Immunol 46, 1214–1223 (2016).

20. N. Fournier et al., FDF03, a novel inhibitory receptor of the immunoglobulin superfamily, is expressed by human dendritic and myeloid cells. J Immunol 165, 1197–1209 (2000).

21. R. A. Wilcox et al., Provision of antigen and CD137 signaling breaks immunological ignorance, promoting regression of poorly immunogenic tumors. J Clin Invest 109, 651–659 (2002).

