## Supplemental figures 1 and 2 for "The in vivo inhibitory function of the MHC-I α3 domain–CD8α interaction"

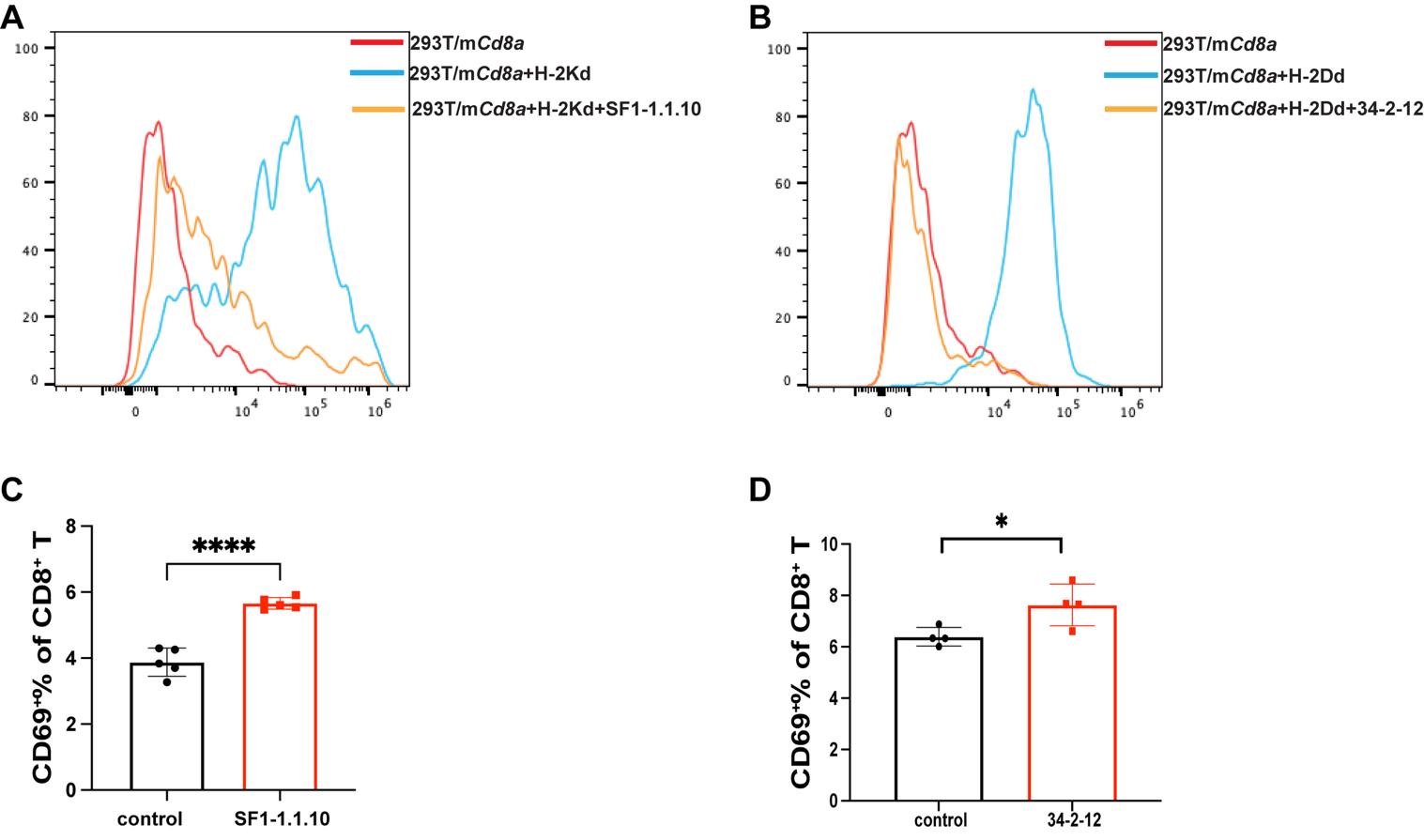
**Fig. S1. Blockade of H-2K^d^ α3 domain/CD8α interaction or H-2D^d^ α3 domain/CD8α interaction could lead to spontaneous activation of peripheral CD8+ T cells**.

(**A**). Flow cytometry analysis of fluorescence-labeled H-2K^d^ tetramer binding to mouse CD8a-transfected 293T cells in the absence or presence of anti-mouse H-2K^d^ α3 domain mAb (clone: SF1-1.1.10).

(**B**), Flow cytometry analysis of fluorescence-labeled H-2D^d^ tetramer binding to mouse CD8a-transfected 293T cells in the absence or presence of anti-mouse H-2d^d^ α3 domain mAb (clone: 34-2-12).

(**C**), One day after control or SF1-1.1.10 antibody treatment, CD69+ % in the CD8+ T cells in the lymph node was detected by flow cytometry.

(**D**), One day after control or 34-2-12 antibody treatment, CD69+ % in the CD8+ T cells in the lymph node was detected by flow cytometry.

Data are presented as mean ± SD. n = 5 mice per group in (C) and n = 4 mice per group in (D). Statistical significance was determined using an unpaired Student’s *t* test. **P* < 0.05; *****P* < 0.0001.

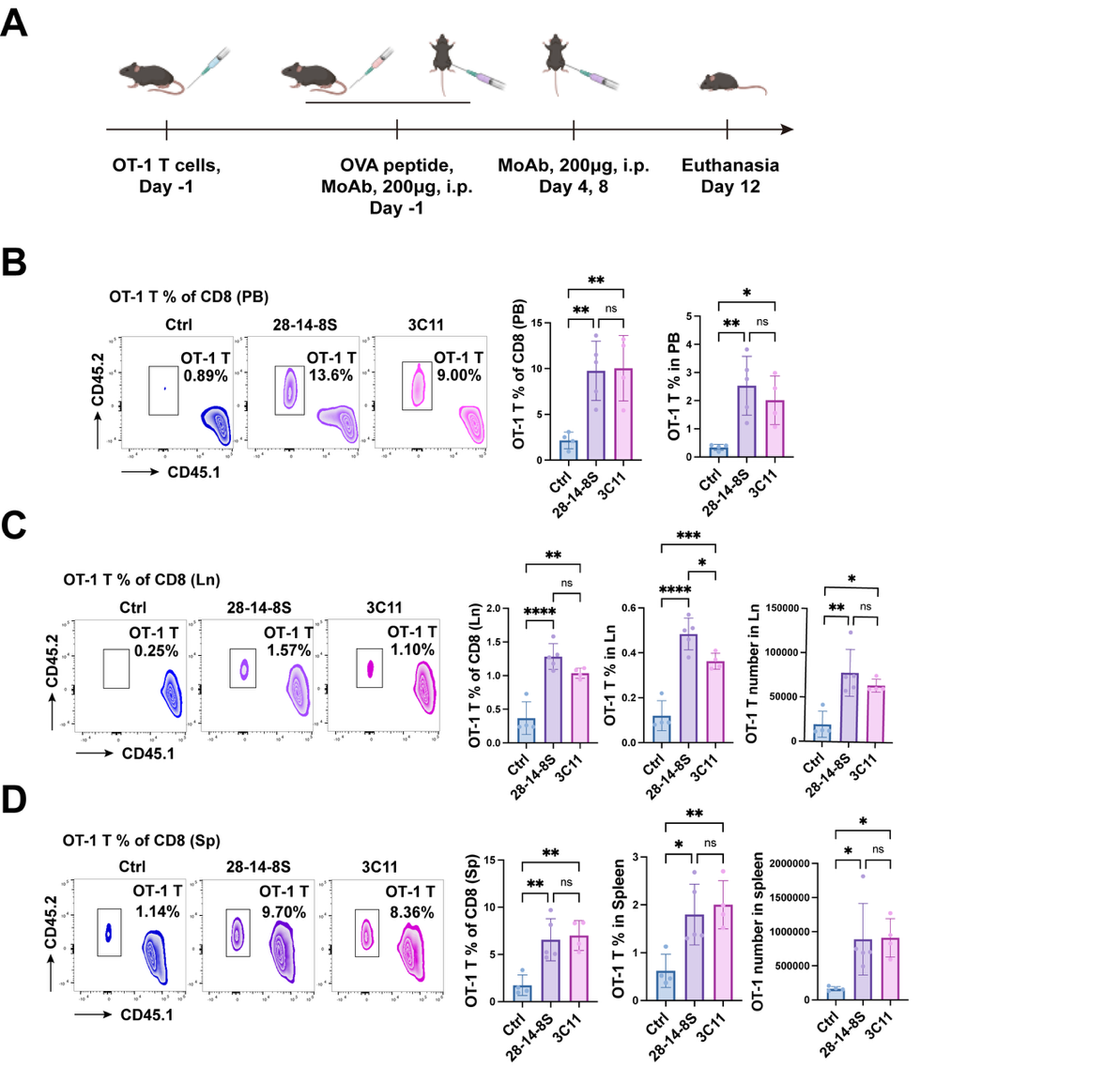
 **Fig. S2. Blockade of either the H-2D^b^ α3 domain–CD8α interaction or the H-2K^b^ α3 domain–CD8α interaction enhances antigen-induced CD8^+^ T cell responses systemically in the blood, lymph nodes, and spleen.**

**(A)** Schematic of the experimental design. Purified OT-I CD8^+^ T cells (CD45.2) were adoptively transferred into congenic CD45.1 C57BL/6 mice.

**(B–D)** On day 12, the frequency and absolute number of OT-I cells (gated as CD45.2^+^CD45.1^−^CD8^+^) in peripheral blood mononuclear cells (PBMCs) (B), lymph nodes (C), and spleen (D) were analyzed by flow cytometry.

Data are presented as mean ± SD. n = 4–5 mice per group. Statistical significance was determined by one-way ANOVA. *P < 0.05; **P < 0.01; ***P < 0.001; ****P < 0.0001; ns, not significant.
